# No evidence of fetal defects or anti-syncytin-1 antibody induction following COVID-19 mRNA vaccination

**DOI:** 10.1101/2021.12.07.471539

**Authors:** Alice Lu-Culligan, Alexandra Tabachnikova, Maria Tokuyama, Hannah J. Lee, Carolina Lucas, Valter Silva Monteiro, M. Catherine Muenker, Subhasis Mohanty, Jiefang Huang, Insoo Kang, Charles Dela Cruz, Shelli Farhadian, Melissa Campbell, Inci Yildirim, Albert C. Shaw, Albert I. Ko, Saad B. Omer, Akiko Iwasaki

## Abstract

The impact of coronavirus disease 2019 (COVID-19) mRNA vaccination on pregnancy and fertility has become a major topic of public interest. We investigated two of the most widely propagated claims to determine 1) whether COVID-19 mRNA vaccination of mice during early pregnancy is associated with an increased incidence of birth defects or growth abnormalities, and 2) whether COVID-19 mRNA-vaccinated human volunteers exhibit elevated levels of antibodies to the human placental protein syncytin-1. Using a mouse model, we found that intramuscular COVID-19 mRNA vaccination during early pregnancy at gestational age E7.5 did not lead to differences in fetal size by crown-rump length or weight at term, nor did we observe any gross birth defects. In contrast, injection of the TLR3 agonist and double-stranded RNA mimic polyinosinic-polycytidylic acid, or poly(I:C), impacted growth *in utero* leading to reduced fetal size. No overt maternal illness following either vaccination or poly(I:C) exposure was observed. We also found that term fetuses from vaccinated murine pregnancies exhibit high circulating levels of anti-Spike and anti-RBD antibodies to SARS-CoV-2 consistent with maternal antibody status, indicating transplacental transfer. Finally, we did not detect increased levels of circulating anti-syncytin-1 antibodies in a cohort of COVID-19 vaccinated adults compared to unvaccinated adults by ELISA. Our findings contradict popular claims associating COVID-19 mRNA vaccination with infertility and adverse neonatal outcomes.

## Introduction

Pregnant women are at increased risk for severe coronavirus disease 2019 (COVID-19) with higher rates of hospitalization, intensive care, and death compared to nonpregnant women [1-7]. However, pregnant women were not included in the initial COVID-19 vaccine trials, leading to a lack of data on vaccine-associated benefits or adverse events in this population [8]. While the first COVID-19 vaccine trials in pregnant women are now underway, over 158,000 women in the United States have self-identified to the Centers for Disease Control and Prevention (CDC) v-safe COVID-19 Pregnancy Registry as having received some form of the vaccine during pregnancy. The mRNA vaccines, Pfizer-BioNTech’s BNT162B2 and Moderna’s mRNA-1273, represent the majority of these immunization events as they were the first to receive emergency use authorization in the United States. Preliminary studies of these data found no serious safety concerns to maternal or fetal health associated with mRNA vaccination but were limited in their assessment of vaccination events during the first and second trimesters due to ongoing pregnancies [9]. These findings from the CDC are notably consistent with the conclusions from other studies supporting the safety of COVID-19 mRNA vaccination during pregnancy [10,11].

Even as more data accumulate supporting the safety of the mRNA COVID-19 vaccines in pregnant and nonpregnant individuals alike [12,13], public perception of the risk surrounding vaccination has been impacted by the rapid proliferation of numerous theories that lack confirmatory data or overtly misrepresent the current evidence.

Women are more likely than men to be vaccine hesitant [14], and a number of these unconfirmed speculations target women’s health issues specifically, spreading fear about pregnancy, breastfeeding, and fertility post-vaccination. Concerns about safety and efficacy remain the most highly cited reasons for COVID-19 vaccine hesitancy in reproductive age and pregnant women [15-17]. Meanwhile, public health recommendations for pregnant women in particular continue to evolve with more data, further complicating the process of vaccine acceptance [18].

Concerns about the impact of mRNA COVID-19 vaccination on future fertility are a major source of vaccine hesitancy in non-pregnant women. One of the most frequently noted concerns is that maternal antibodies generated against the severe acute respiratory syndrome coronavirus 2 (SARS-CoV-2) spike protein in response to vaccination could result in cross-reactivity to the retrovirus-derived placental protein, syncytin-1 [19]. Syncytin-1 is encoded by the human endogenous retrovirus W (*HERV-W*) gene and is involved in trophoblast fusion during placental formation [20]. There is limited homology between syncytin-1 and spike protein of SARS-CoV-2, and this is likely not sufficient to mediate cross-reactivity of vaccine-induced anti-spike antibodies to syncytin-1. Furthermore, many women at this stage in the pandemic have conceived following both COVID-19 infection and COVID-19 vaccination, with no reports of reduced fertility. Despite the absence of supporting evidence [21,22], the link between the mRNA vaccines and infertility persists, undermining mass vaccination campaigns worldwide.

Amidst public pressure and media scrutiny surrounding the potential risks of vaccination during pregnancy, the beneficial effects of vaccination on the fetus *in utero* have received far less attention. Recent evidence has linked COVID-19 vaccination during the third trimester with improved maternal and neonatal outcomes[23]. Following natural infection with SARS-CoV-2, maternal antibodies against SARS-CoV-2 proteins readily cross the placenta[24,25]. Likewise, recent studies have found antibodies against the Spike and RBD domains of SARS-CoV-2 in human infant cord blood samples following vaccination during pregnancy, suggesting vaccine-induced protection by maternal antibodies is conferred across the maternal-fetal interface [26-29]. Maternal transfer of antibody-mediated protection following vaccination against SARS-CoV-2 to the infant can additionally occur via breastmilk in nursing women [30-33].

Mouse models permit an investigation of vaccine effects at defined points of organogenesis at the earliest periods of pregnancy, timepoints at which many women do not yet know they are pregnant. Vaccine doses can also be administered to mice at concentrations many times greater than that used in humans to maximize the probability of eliciting and observing any potential teratogenic effects. In this study, we investigated the impact of mRNA-1273 vaccination during early pregnancy using a mouse model to screen for birth defects and quantified circulating antibodies in mother and fetus at the end of gestation. Finally, using human samples, we analyzed the effect of mRNA-1273 (Moderna) and BNT162B2 (Pfizer-BioNTech) vaccination on circulating anti-syncytin-1 antibody levels to address infertility speculation.

## Results

### Vaccination of pregnant mice during early pregnancy with mRNA-1273 does not lead to fetal birth defects or differences in fetal size

To determine whether COVID-19 mRNA vaccination during early pregnancy leads to birth defects in mice, pregnant dams were subjected to intramuscular (i.m.) injection of 2 µg mRNA-1273 at E7.5 and fetuses were harvested at term but prior to birth at E18.5 to assess phenotypes. We chose this early timepoint of pregnancy for vaccination, as previous studies have demonstrated particular vulnerability to the development of fetal defects by stimulating maternal innate immune responses during this gestational period [34-39]. We injected a very large dose of mRNA vaccine, 2 µg mRNA-1273 in an average mouse weighing 25 g corresponding to over 50 times the µg vaccine per g weight administered to humans, to ensure we detect impact of vaccine on the developing fetus, if any. We did not observe any birth defects in either vaccine-exposed or PBS-treated control litters (Figure 1A, B). Fetal length and weight measured directly prior to birth at E18.5, or term prior to parturition, were also not impacted by maternal vaccination during early gestation (Figure 1C, D).

**Figure 1.**
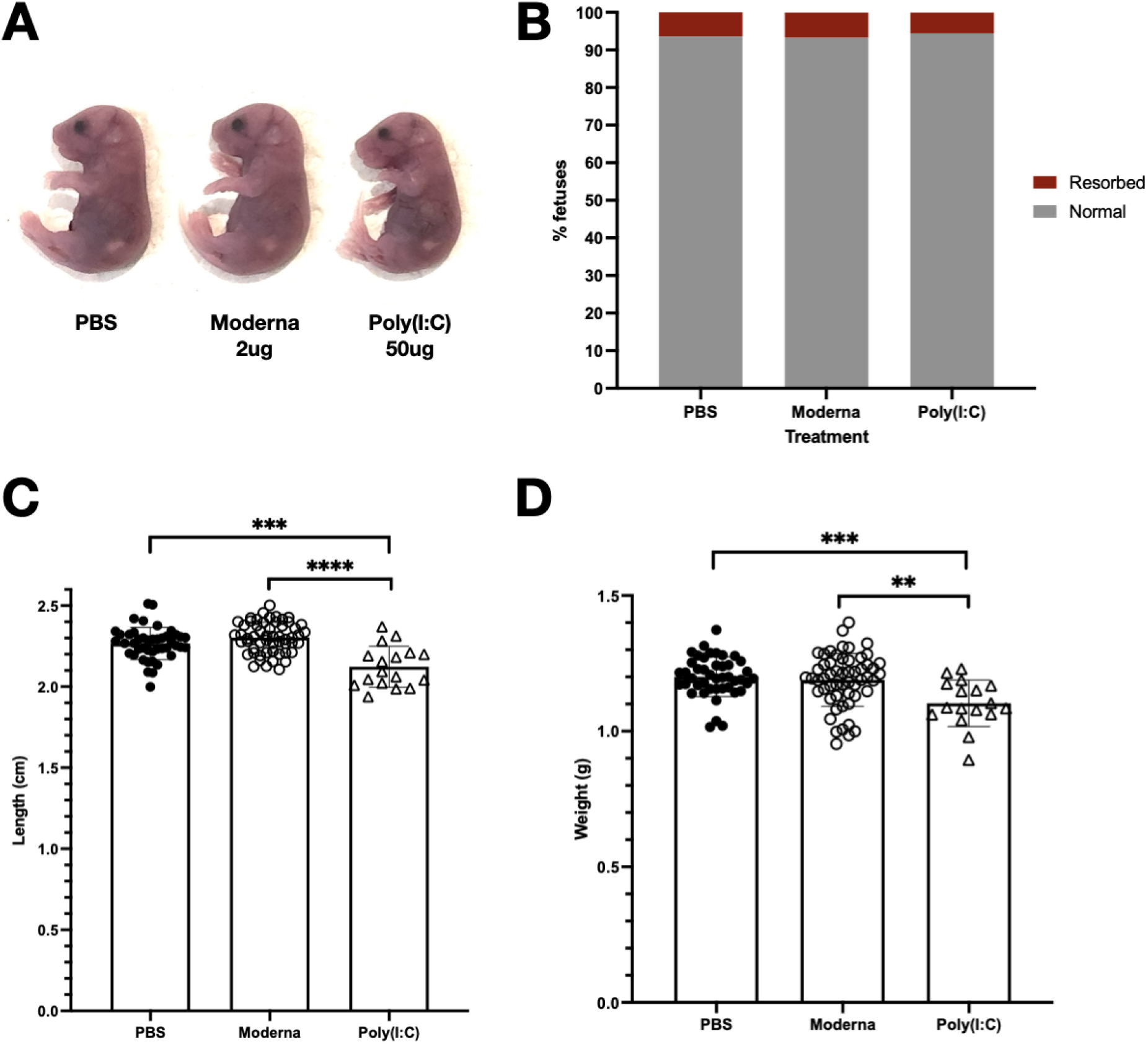
Maternal vaccination with Moderna mRNA-1273 at E7.5 in early pregnancy does not impact fetal viability and growth at the end of gestation. Pregnant dams were injected intramuscularly with 50ul of PBS (n=6 litters, 44 fetuses), 2 µg Moderna vaccine (n=7 litters, 55 fetuses), or 50 µg poly(I:C) (n=3 litters, 17 fetuses) at E7.5 and fetuses were harvested at E18.5. (A) Normal phenotypes observed in fetuses from PBS-treated and Moderna-vaccinated pregnancies. (B) Fetal resorption rates in PBS-treated, Moderna-vaccinated, and Poly(I:C)-treated pregnancies. (C) Crown-rump length and (D) weight of dissected fetuses. Shapiro-Wilk normality tests were used to confirm Gaussian distribution of fetal weights and crown-rump lengths. Brown-Forsythe and Welch ANOVA tests were used to calculate statistical significance (*: p ≤ 0.05; **: p ≤ 0.01; ***: p ≤ 0.001; ****: p ≤ 0.0001).

Administration of poly(I:C), a double-stranded RNA mimic and potent TLR3 agonist, induces maternal immune activation and results in fetal growth restriction and fetal demise in rodents [39-41]. Compared to mRNA-vaccinated and control groups, maternal injection i.m. with a high dose of 50 µg poly(I:C) at E7.5 resulted in decreased fetal crown-rump length and weight at E18.5. No other birth defects were seen following i.m. delivery of poly(I:C).

## Anti-Spike antibodies are detected in fetal circulation following maternal vaccination

In order to test whether vaccine-induced antibodies against SARS-CoV-2 can cross the maternal-fetal interface, we quantified anti-Spike and anti-RBD antibodies in the fetal sera by ELISA following vaccination of dams during early pregnancy. Pregnant dams were vaccinated i.m. with mRNA-1273 at E7.5 and both maternal and fetal sera were analyzed 12 days post-vaccination at E18.5, prior to birth. Both maternal and fetal sera from vaccinated pregnancies contained high levels of circulating antibodies against SARS-CoV-2 Spike and RBD by ELISA as compared sera from PBS-injected mice (Figure 2).

**Figure 2.**
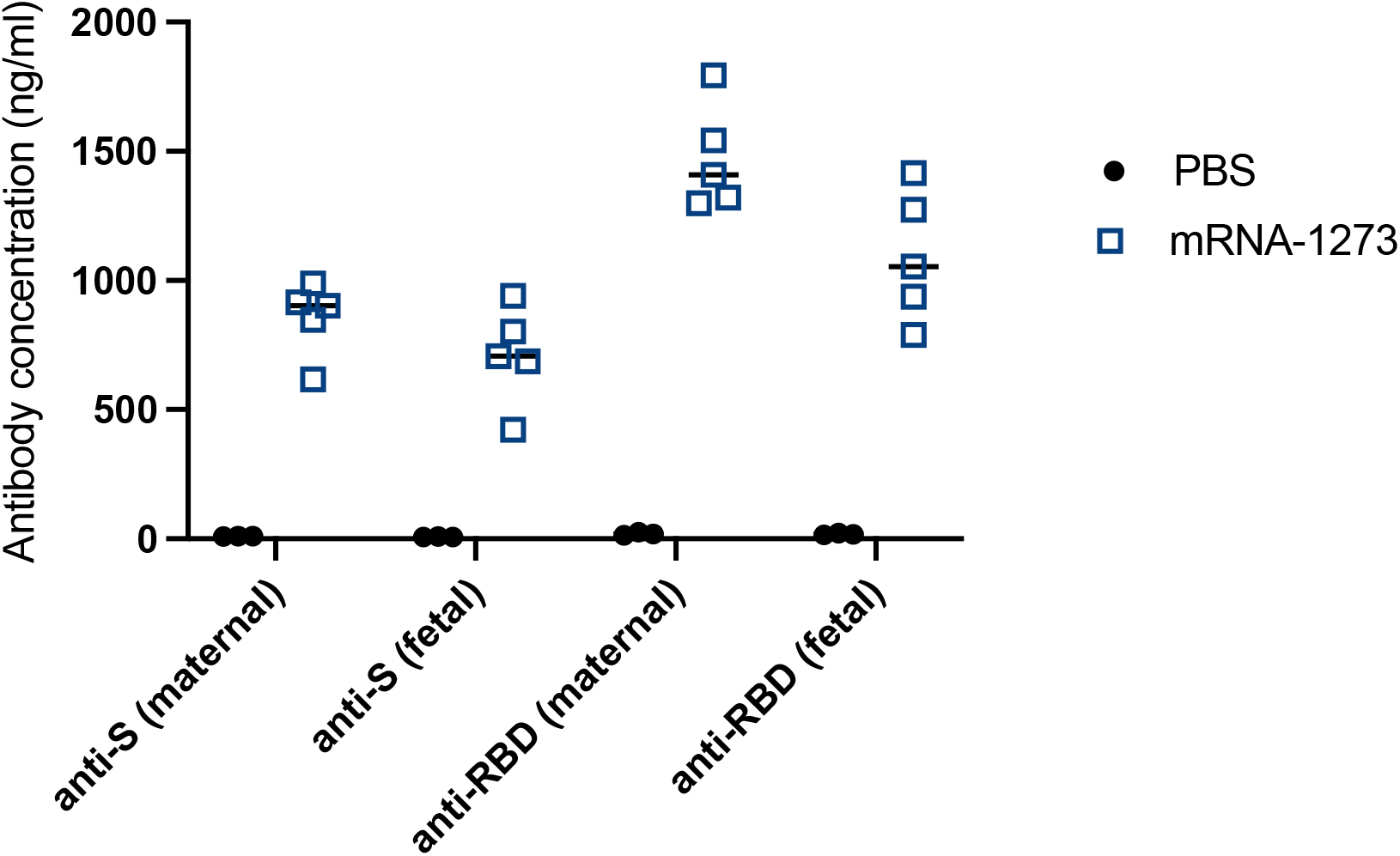
Maternal vaccination against SARS-CoV-2 induces an antibody response that crosses the maternal-interface and is detectable in fetal sera. Pregnant dams were injected intramuscularly with 50µl of PBS (n=3 litters) or 2µg Moderna vaccine (n=5 litters) at E7.5. Maternal serum and pooled fetal serum from each litter were collected at E18.5, 12 days post-treatment. Anti-Spike (anti-S) and anti-Receptor-Binding Domain (anti-RBD) levels were measured by ELISA and antibody concentration was calculated based on a standard curve.

## COVID-19 vaccination does not induce anti-syncytin-1 autoimmunity/antibodies in vaccinated people

We quantified levels of anti-syncytin-1 antibodies in human sera from a cohort of unvaccinated and vaccinated adult volunteers [42] to determine whether vaccination status is associated with anti-syncytin-1 antibodies. To calculate the concentration of anti-syncytin-1 antibodies, we generated a standard curve using a commercially available monoclonal antibody (Supplementary Figure 1).

We compared the levels in these groups to a previously collected cohort of patients with systemic lupus erythematosus (SLE) [43]. SLE is a complex disease with variable presentation that is associated with ERV dysregulation and elevation in anti-ERV envelope antibodies[43]. We detected two SLE patients with highly elevated anti-syncytin-1 antibody levels, while the remainder were negative (Figure 3, Supplementary Figure 1).

**Figure 3.**
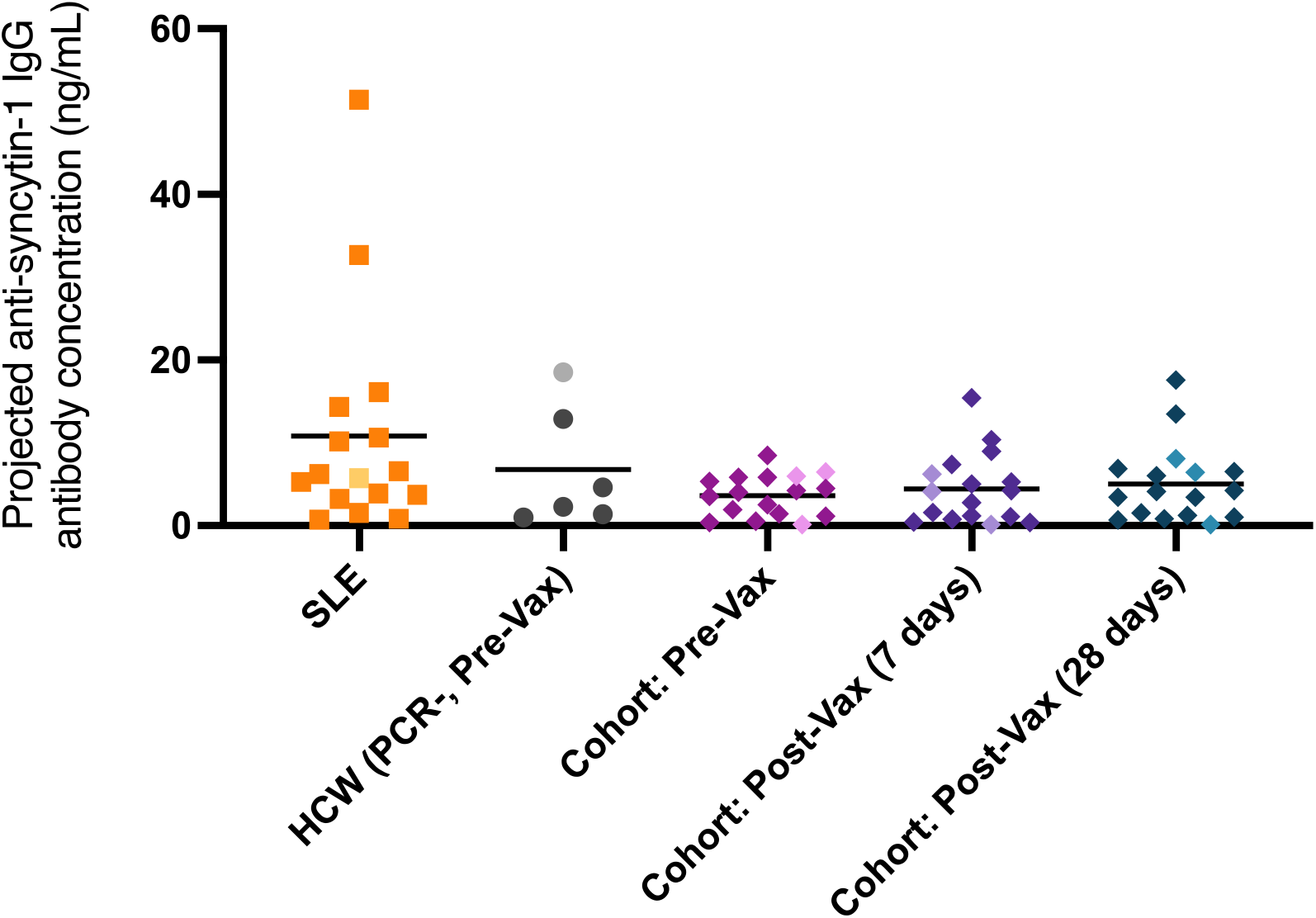
mRNA vaccination against COVID-19 is not associated with increased levels of circulating anti-syncytin-1/HERV-W IgG antibodies in humans. Plasma reactivity to syncytin-1 protein was assessed using ELISA. SLE samples (n=16) and uninfected, unvaccinated HCW controls (SARS-COV-2 PCR negative, pre-vaccination; n=6) are shown alongside a cohort of matched samples from individuals pre-vaccination, 7 days post-second vaccine dose, and 28 days post-second vaccine dose of mRNA-1273 or BNT162B2 (n=17). Each dot represents a single individual and male subjects are lightened in color. Horizontal bars represent mean values. Statistical significance was assessed using nonparametric Kruskal-Wallis tests and Mann-Whitney tests. No groups were significantly elevated. Antibody concentrations were calculated based on a standard curve generated using a monoclonal antibody against syncytin-1.

Both unvaccinated and COVID-19 mRNA vaccinated groups had lower levels of anti-syncytin-1 antibodies than the SLE patients identified with higher anti-syncytin-1 levels. Unvaccinated healthcare workers with no known history of SARS-CoV-2 positivity by PCR and ELISA were used as a control group, exhibiting anti-syncytin-1 levels comparable to the antibody-negative SLE patients.

We then compared serum levels of anti-syncytin-1 antibodies in matched samples from individuals at three timepoints: 1) pre-vaccination, 2) 7 days post-vaccination (second dose), and 3) 28 days post-vaccination (second dose) with either mRNA-1273 or BNT162B2. The two latter timepoints correspond with the highest titers of anti-SARS-CoV-2 antibodies, peaking at 7 days post-vaccination with the second dose [42].

Anti-syncytin-1 IgG levels in COVID-19 vaccinated adults at 7 and 28 days post-vaccination with a second mRNA dose were unchanged from pre-vaccination levels of anti-syncytin-1 in SARS-CoV-2-negative and comparable to unvaccinated control groups (Figure 3). We also compared levels of anti-syncytin-1 IgG during acute COVID-19 disease in pregnant and non-pregnant patients. We did not detect significant levels of anti-syncytin-1 antibodies during the acute phase of SARS-CoV-2 infection in either group (Supplementary Figure 2).

## Discussion

Public fear surrounding the consequences of vaccination on pregnancy and fertility remains an ongoing source of vaccine hesitancy during the COVID-19 pandemic. Here, we demonstrate in a mouse model that intramuscular injection of the mRNA-1273 vaccine during early pregnancy does not induce birth defects and does not lead to differences in fetal size at birth. These findings are consistent with CDC data in humans that found no congenital anomalies following vaccination in the first trimester or periconception periods, although these numbers were limited [9]. More extensive data on vaccination during the second and third trimesters have similarly shown a high safety profile with no increase in adverse outcomes [9]. Our results did not uncover overt fetal defects in mice exposed to early pregnancy vaccination with an mRNA-1273 dose approximately 50 times greater than that used in humans. The vaccine dosage used in these murine studies (2 µg total, or 0.08 µg mRNA-1273 per gram weight in a 25 g mouse) are over 50 times the dose by weight used in humans (100 µg regardless of weight, or 0.00125 µg mRNA-1273 per gram weight for average individual of 80 kg in the U.S.). We specifically chose this high dose to test whether COVID-19 mRNA vaccine has any potential deleterious effects on pregnancy. Our data demonstrated no negative impact of the vaccine on the pregnant mother or the developing fetus.

Intramuscular administration of poly(I:C), a synthetic double-stranded RNA, to pregnant dams at the same timepoint in early pregnancy also did not result in birth defects. However, poly(I:C)-treated pregnancies were associated with decreased fetal size by crown-rump length and weight at the end of gestation.

The observed differences in fetal outcomes for mRNA vaccine-treated pregnancies and poly(I:C)-treated pregnancies may be attributed to key differences in the immune response induced. The mRNA vaccines, including mRNA-1273, are designed to exhibit reduced immunogenicity with targeted modifications such as N1-methylpseudouridine (m1Ψ) substitutions that decrease TLR3 activation [44]. In contrast, poly(I:C) leads to the robust activation of innate immune sensors such as TLR3. It is well established that poly(I:C) or viral infection during early pregnancy in mice leads to fetal growth restriction and demise [40,45,46]. In contrast, even at high doses, mRNA vaccine resulted in no negative impact on the developing fetus. These findings further suggest that the immune response generated after mRNA vaccination is safer in pregnancy than the immune response to SARS-CoV-2 infection, which is known to elicit a robust inflammatory signature at the maternal-fetal interface [47].

Our data also demonstrated that pregnant dams vaccinated during early pregnancy, prior to the establishment of fetal circulation, subsequently confer protective antibodies to the *in utero* fetus up to the time of birth. These findings are consistent with human studies showing the presence of SARS-CoV-2 vaccine-induced antibodies in cord blood following delivery and currently limited data on first trimester vaccination [26-29].

Recent studies suggest that the timing of vaccination during pregnancy may be important for the efficiency of transplacental transfer of antibodies from mother to fetus, with some pointing to a beneficial impact of vaccination earlier in pregnancy during the second or early third trimester compared to later in the third trimester [48-53]. Though many women have now received COVID-19 vaccinations in the first trimester, few studies had previously been able to focus on immunization during these early timepoints. Given this interest, continuation of this work is thus necessary to comprehensively investigate both the safety and potential of earlier vaccination schedules.

Limitations of our mouse model of vaccination include the use of a single dose of mRNA-1273 during pregnancy whereas the full vaccine schedule in humans is two doses approximately 28 days apart, longer than the murine gestation period. While this study provides the first step in establishing safety for early vaccine approaches in pregnancy, we surveyed litters only for overt birth defects and size at E18.5. Findings of this work await validation with results from the ongoing human studies of vaccination during the first trimester. Our study does not capture potential effects of mRNA vaccination timing during late gestation.

Finally, using human data we demonstrated that circulating anti-syncytin-1 antibodies do not rise following COVID-19 mRNA vaccination with either mRNA-1273 or BNT162B2. These findings support the mounting evidence that a syncytin-1-based mechanism of infertility is not supported by scientific observations [19,21]. This study is limited as we only interrogated the presence of anti-syncytin-1 in a total of 51 subjects. Larger studies are warranted to further assess anti-syncytin-1 antibody status in women who receive the mRNA vaccines.

Millions of women and pregnancies continue to be impacted by the ongoing COVID-19 pandemic and by vaccine hesitancy. In the absence of complete clinical trial data, many women are choosing to vaccinate during pregnancy after weighing the risks and benefits of their situation. Thus, filling in gaps in knowledge, particularly surrounding vaccination in early pregnancy, and combatting misinformation are more imperative than ever. This study thus provides a reassuring view of the safety and protection provided following COVID-19 mRNA vaccination during early pregnancy within a mouse model. We demonstrate the absence of anti-syncytin-1 antibodies in our cohort of vaccinated adults, discrediting one of the most widespread infertility myths surrounding COVID-19 vaccination. Future work must be expanded to examine the impact of vaccination at all gestational ages and continue to provide data that can address public concerns about vaccine safety in all populations.

## Acknowledgments

We thank all patients and healthcare workers who generously participated in this study. We also thank the Yale IMPACT Team for their contribution to this research. A.I. is an Investigator of the Howard Hughes Medical Institute.

## Competing Interests

Authors have no competing interests to declare.

## Methods

### Timed mating and injections

C57BL/6J mice were mated overnight and females were checked for the presence of seminal plugs each morning, designated E0.5. On E7.5, pregnant mice were anesthetized and subjected to a single intramuscular (i.m.) injection into the thigh muscle of the hind limb with 50ul volume of PBS, 2 µg of mRNA-1273, or 50 µg poly(I:C). Vaccigrade HMW poly(I:C) (Invivogen #vac-pic) was prepared at 1 mg/ml at room temperature and stored at -20C, then thawed to room temperature prior to injection.

### Harvest of fetuses, fetal serum collection, and fetal measurements

On E18.5, prior to birth, pregnant dams and fetuses were harvested. Fetuses were measured, weighed, and assessed for birth defects such as missing eye(s) and neural tube defects. Fetal blood was collected and allowed to clot for 1 hour at room temperature before two rounds of centrifugation at 10000 x g for 10 minutes at 4C. Serum was collected and stored at -80C.

All statistical analyses comparing fetal viability and growth measurements were performed with GraphPad Prism 8.4.3 software. Before assessing the statistical significance, Shapiro-Wilk normality test was used to confirm Gaussian distribution of the fetal weights and crown-rump lengths. Afterwards, the data were analyzed with Brown-Forsythe and Welch ANOVA tests for multiple comparisons between the PBS, Poly(I:C), and Moderna treatments (∗: p ≤ 0.05; ∗∗: p ≤ 0.01; ∗∗∗: p ≤ 0.001; ∗∗∗∗: p ≤ 0.0001).

### Human plasma collection and cohort selection

Plasma was collected from healthcare worker (HCW) volunteers who received the mRNA vaccine (Moderna mRNA-1273 or Pfizer-BioNTech BNT162B2) between November 2020 and January 2021 as approved by the Yale Human Research Protection Program Institutional Review Board (IRB Protocol ID 2000028924). None of the participants experienced serious adverse effects after vaccination. HCWs were followed serially post-vaccination, and samples were collected at baseline (prior to vaccination), and 7- and 28-days post second vaccination dose. Blood acquisition was performed and recorded by a separate team. Female HCWs age 55 and under were selected, along with 3 male controls. Plasma samples from patients experiencing acute infection with SARS-CoV-2 ranging from asymptomatic to severe disease, including six pregnant patients, were selected from the previously described Yale IMPACT study, approved by the Yale Human Research Protection Program Institutional Review Board (FWA00002571, IRB Protocol ID 2000027690)[54]. Matched uninfected, unvaccinated HCW were also analyzed.

Finally, plasma from previously described[43] patients with systemic lupus erythematosus (SLE), was used as a positive control. Female patients under the age of 60 were selected, as well as one male SLE patient. SLE patients were recruited from the rheumatology clinic of Yale School of Medicine and Yale New Haven hospital in accordance with a protocol approved by the institutional review committee of Yale University (#0303025105). The diagnosis of SLE was established according to the 1997 update of the 1982 revised American College of Rheumatology criteria [55] [56]. After obtaining informed consent, peripheral blood was collected in EDTA tubes from human subjects, and plasma was extracted upon centrifugation. Plasma were stored at −80°C. All patients had SLE according to the American College of Rheumatology criteria for classification of SLE.

### Isolation of human plasma

Whole blood from HCW was collected in heparinized CPT blood vacutainers (BD #BDAM362780) and kept on gentle agitation until processing. All blood was processed on the day of collection in a single step standardized method. Plasma samples were collected after centrifugation of whole blood at 600 x g for 20 minutes at room temperature without brake. The undiluted plasma was transferred to 15-ml polypropylene conical tubes, and aliquoted and stored at −80 °C for subsequent analysis.

### Anti-HERV-W/Syncytin-1 antibody and ELISAs

96-well MaxiSorp plates (Thermo Scientific #442404) were coated with 20 ng per well of human Syncytin-1 recombinant protein (Abnova #H00030816-Q01) in PBS and were incubated overnight at 4°C. The coating buffer was removed, and plates were blocked overnight at 4°C with 250 μl of blocking solution (PBS with 0.1% Tween-20, 3% milk powder). Plasma was diluted 1:800 in dilution solution (PBS with 0.1% Tween-20, 1% milk powder) and 100 μl of diluted plasma was added for one hour at room temperature. Mouse Anti-HERV-W monoclonal antibody (Abnova #H00030816-M06) was serially diluted to generate a standard curve. All samples were plated in duplicate. Plates were washed three times with PBS-T (PBS with 0.1% Tween-20) and 50 μl of HRP anti-Human IgG Antibody (GenScript #A00166) diluted 1:5000 in dilution solution were added to each well. 50 ul of HRP anti-Mouse IgG1 Antibody (Southern Biotech #1070-05) diluted 1:3000 in dilution solution were added to each standard well. After one hour of incubation at room temperature, plates were washed six times with PBS-T. Plates were developed with 50 μl of TMB Substrate (Invitrogen #00-4201-56) and the reaction was stopped after 10 minutes by the addition of 50 μl 2 N sulfuric acid. Plates were then read at a wavelength of 450 nm and 570nm. To fit the standard curve, mean absorbance (OD450nm) was plotted known antibody concentration to generate a standard curve. Best fit was determined using asymmetrical sigmoidal five-parameter least-squares fit in GraphPad Prism 9.2.0. Projected antibody concentrations were interpolated using this fit.

Statistical significance (p) was determined using nonparametric Kruskal-Wallis test followed by Dunn’s multiple comparisons test for matched samples across vaccination time points (Figure 3), and by nonparametric Mann-Whitney test to compare each individual cohort to uninfected, unvaccinated HCW controls or to SLE patients (Figure 3, Supplementary Figure 2). All analyses were two-tailed and carried out in GraphPad Prism 9.2.0.

**Supplemental Figure 1.**
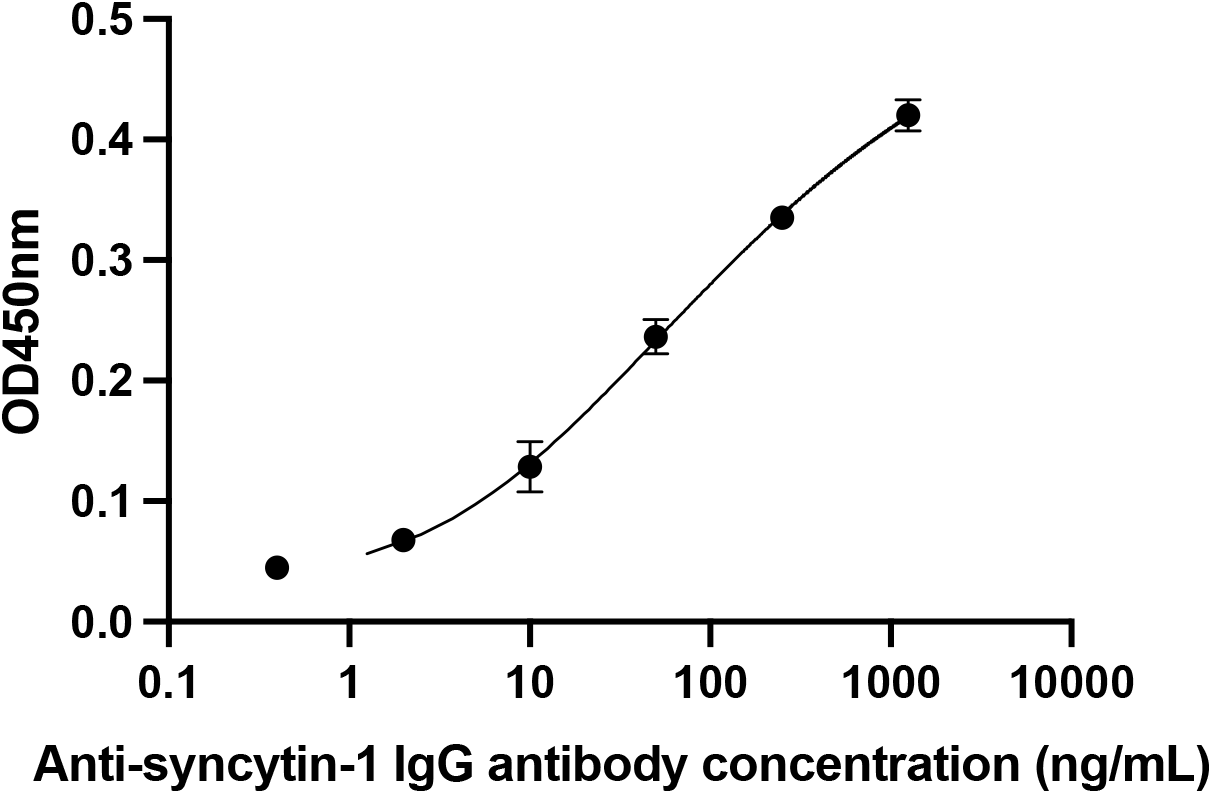
Standard curve for anti-syncytin-1/HERV-W monoclonal antibody. Mean absorbance (OD450nm) plotted against serial dilutions of monoclonal IgG antibody against syncytin-1 to generate a standard curve. Best fit was determined using asymmetrical sigmoidal five-parameter least-squares fit. Projected antibody concentrations were interpolated using this fit. Showing one of six replicates. R-squared = 0.9996, Sum of squares = 6.572e-005.

**Supplemental Figure 2.**
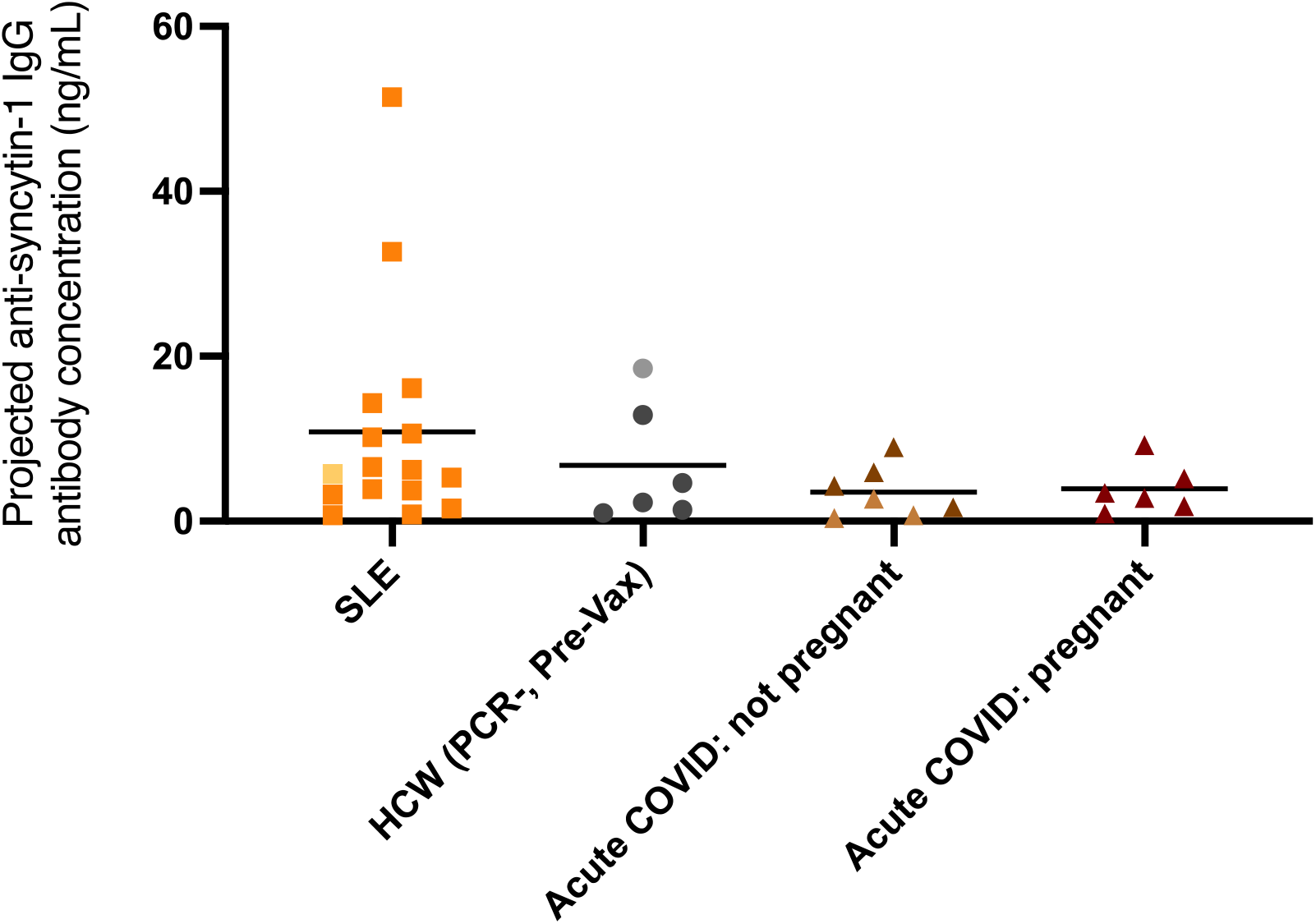
Acute COVID-19 disease is not associated with increased levels of circulating anti-syncytin-1/HERV-W antibodies in humans. Plasma reactivity to syncytin-1 protein was assessed by ELISA in SLE samples (n=16), unvaccinated HCW samples (n=6), non-pregnant patients with acute COVID-19 disease (n=7), and pregnant patients with acute COVID-19 disease (n=6). Each dot represents a single individual. Male subjects are lightened in color. Horizontal bars represent mean values. Statistical significance was assessed using nonparametric Mann-Whitney tests. No groups were significantly elevated as compared to HCW controls.

## Notes

### Competing Interest Statement

The authors have declared no competing interest.

